# Unified understanding of nonparametric causality detection in time series

**DOI:** 10.1101/2023.04.20.537743

**Authors:** Yutaka Osada, Masayuki Ushio, Michio Kondoh

## Abstract

Most complex systems in the real world are driven by multiple interactions among components. Identifying these interactions is critical for understanding, forecasting, and controlling system-level phenomena. Transfer entropy (TE) and convergent cross mapping (CCM) are two widely-used approaches for nonparametric causality detection based on time-series data. However, the theoretical relationship between TE and CCM has not been formally explained. Here, we provide a theoretical formulation that links TE and CCM in an information-theoretic framework, showing that they have different definitions of causal influence. Furthermore, from the formulation, we propose a novel nonparametric causality test, named unified information-theoretic causality (UIC), which has lower data requirements than those of TE (due to robustness to noise) and lower false-positive rates than those of CCM. Numerical experiments confirmed that UIC outperforms TE and CCM in terms of the performance of causality detection for both linear and nonlinear dynamical systems. Finally, we demonstrate the practical importance of the conditional test based on UIC using numerical simulations with indirect causality and empirical data for microbial communities in irrigated water samples from rice fields. Our results contribute to a unified understanding of nonparametric causality tests in time series and the accurate reconstruction of complex real-world systems.

## I. INTRODUCTION

Detection of cause–effect relationships among variables, events or objects is a fundamental topic in both natural and social sciences. Most attempts at causality detection are easily disturbed by confounding factors [1] and nonlinear dynamics [2]. These issues are particularly challenging in large systems where intervention is technically or ethically difficult or even impossible [3]. In recent years, nonparametric tests for time series have attracted attention as promising methods for detecting potential causal relations with the minimal need to intervene in the target system [2, 4]. Time-series data support Granger’s statistical concepts of causality: (1) the cause necessarily occurs before the effect, and (2) the cause contains information about the effect when accounting for confounding variables [5, 6].

Nonparametric causality tests based on time-series data have been proposed from the perspectives of information theory and dynamical systems theory. In this study, we focus on two widely-used methods: transfer entropy (TE) [7] and convergent cross mapping (CCM) [2]. TE is an information-theoretic causality test, known as a nonparametric generalization of Granger causality [8]. TE has a clear mathematical definition of causal influence, which is advantageous for theoretical advances [9, 10] and practical applications [3, 11]. CCM is a non-parametric causality test developed mainly for nonlinear dynamical systems [2, 12]. The most important feature of CCM is the use of time-delay embedding theorems [13], which allow for the detection of causal relations without addressing confounding variables embedded by state space reconstruction [2]. As the performance of these methods can be reversed depending on the system [2, 14], it is essential to clarify the mathematical connection between TE and CCM in order to understand their advantages and disadvantages as causal detection in various systems.

Here, we provide a theoretical formulation that links TE and CCM in an information-theoretic framework. Furthermore, based on the formulation, we propose a novel nonparametric causality test, named unified information-theoretic causality (UIC). UIC incorporates the advantages of both TE and CCM, including (1) a clear mathematical definition of causal influence, (2) noise robustness for causal variables in nonlinear dynamical systems, and (3) efficient computation of statistical significance. Based on the previously developed extension of TE [9], we also introduce a conditional UIC test and apply it to an artificial model system and an empirical microbial system. In the numerical experiments, UIC outperformed TE and CCM for both linear and non-linear dynamical systems, consistent with the theoretical results.

## II. UNIFYING CAUSALITY TESTS

### A. Setting

We consider a multivariate time series with ***x*** = {*x*_*t*_} as the effect variable and ***y*** = {*y*_*t*_} as the cause variable and test whether *y*_*t−p*_ has a causal influence on *x*_*t*_ with time lag *p >* 0. Under generic conditions, *x*_*t*_ can be embedded by *y*_*t−p*_ and 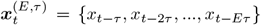 based on the time-delay embedding theorem [13, 15]. Note that, in most real-world cases, the embedding dimensions (*E >* 0) and time interval (*τ >* 0) are not known *a priori* and need to be estimated [16].

### B. TE and CCM

In the above setting, we give the definitions of TE and CCM following their original formulations. Using the joint probability *p*(*x, y*) and the conditional probability *p*(*x*|*y*), TE measures causal influence as the flow of information from a cause variable to an effect variable as follows [7]:

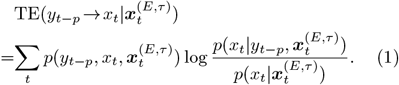

Thus, the causal influence from *y*_*t−p*_ to *x*_*t*_ is tested by rejecting the null hypothesis, 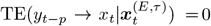. CCM evaluates the predictive performance of cross mapping (i.e., a nonparametric regression model based on the nearest neighbor method) to test for causal influence. Under the assumption of Gaussian noise, the predictive performance *X*(***L, P***) is defined as follows [2]:

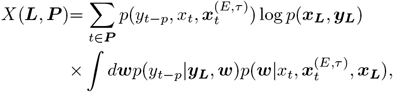

where ***L*** and ***P*** are the sets of time indices for state space reconstruction (i.e., model training) and prediction (i.e., model cross-validation), respectively. ***x***_***L***_ = {*x*_*t*_|*t* ∈ ***L***} and ***y***_***L***_ = {*y*_*t*_|*t* ∈ ***L***}. ***w*** denotes the model parameters (e.g., projection weights). Note that 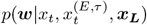 and *p*(*y*_*t−p*_ | ***y***_***L***_, ***w***) describe the process where nearest neighbors are searched from the effect variable and the process where the predictive performance of the cause variable is evaluated based on the searched nearest neighbors, respectively. Sugihara *et al*. [2] measured causal influence by the improvement in predictive performance with increasing training data (so-called “convergence”) as follows:

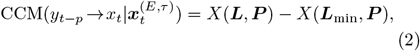

where ***L***_min_ is the set of time indices of training data at the minimum size (minimum library length) [16]. As the training data size approaches zero, we can prove that the expectation of causal influence defined by Eq. (2) is equivalent to that of the following mutual information (see Supplemental Material S1):

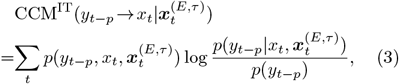

which is considered the information-theoretic definition of the CCM causal influence.

### C. Unified Information-theoretic Causality

Using the information-theoretic definitions, here we clarify the theoretical connection between TE and CCM. We define the following novel measure of causal influence, hereafter referred to as unified information-theoretic causality (UIC):

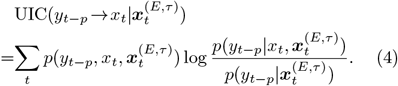

By the Bayes’ rule, *p*(*y*|*x, z*)*p*(*x*|*z*) = *p*(*x*|*y, z*)*p*(*y z*), we can immediately prove the equivalence between TE and UIC from Eqs. (1) and (4). If there are enough data to accurately estimate the conditional probabilities, we will have the same value of causal influence for TE and UIC. It is also obvious from Eqs. (3) and (4) that CCM and UIC (or TE) are equivalent only if *y*_*t−p*_ and 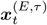 are independent. In general, CCM tends to perform better for nonlinear systems because this independence can be more satisfied due to long-term unpredictability. CCM is more likely to detect false causality than TE [3, 14] because *y*_*t−p*_ may have causal influence on *x*_*t*_ via 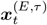 (i.e., the previous states of *x*_*t*_).

In CCM and UIC, ***x*** and ***y*** show complete separation as conditional and predicted variables for conditional probabilities. This variable separation is not trivial for non-parametric causality tests for two reasons. First, it relates to noise robustness because noise of conditional variables has a significant effect on the estimation of causal influence, particularly in nonlinear dynamical systems. As CCM and UIC do not include any cause variables (i.e., *y*_*t−p*_) as conditional variables, they show greater robustness to noise than that of TE. Second, variable separation allows us to efficiently generate surrogate data (see Supplemental Material S3 for details), which are required to evaluate the statistical significance of causal influences [17]. These theoretical expectations are indeed confirmed in the following numerical experiments.

### D. Numerical Experiments

Using artificial model systems, we compared the detection performance of three causality tests. Synthetic timeseries were generated by numerical simulations from two dynamical systems with varying noise: a nonlinear logistic model and linear vector autoregression (VAR) model (Supplemental Material S5). The performance of causality tests was evaluated by the area under the receiver-operator characteristic curve (AUC). We found that UIC outperforms TE and CCM for both nonlinear and linear systems (Fig. 1). The performance of CCM was similar to that of UIC for the nonlinear system (Fig. 1a). However, for the linear system, with an increasing number of time points, the improvement in performance was slower for CCM than for TE and UIC (Fig. 1b) because false positives do not decrease in CCM; even with an infinite number of observations, CCM did not necessarily detect true causality (i.e., a lack of statistical consistency). Consistent with its theoretical expectation, TE was quite sensitive to the noise levels of causal variables for the nonlinear dynamical system and thus requires a considerable number of time points for robust causality detection (Fig. 1a).

**FIG. 1.**
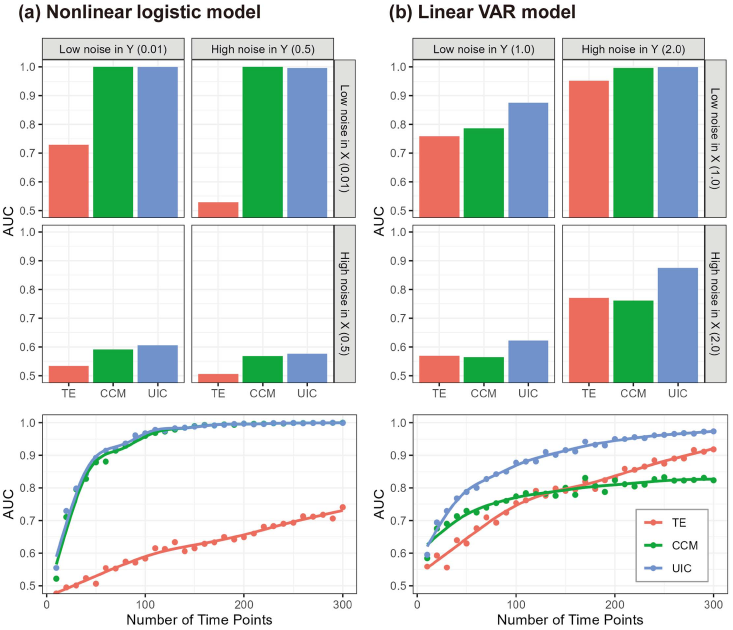
Simulation experiments to compare the detection performance of transfer entropy (TE, red), convergent cross mapping (CCM, green), and unified information-theoretic causality (UIC, blue) in (a) nonlinear logistic model systems and (b) linear VAR model systems. In the upper panels, areas under the receiver-operator characteristic curve (AUC) are compared for different noise levels of effect (*X*) and cause (*Y*) variables. In the lower panels, we examine the relation between data requirements (i.e., the number of time points in time-series data) and AUC at the same noise level. Lines are obtained by fitting simulation results (points) by generalized additive models.

## III. DEVELOPMENT OF CONDITIONAL CAUSALITY TESTS

In most large real-world systems, the causal effect between two variables can be detected by causality tests even when there are no direct relations between them. There are two possible scenarios for the misidentification of causal effects. In the first scenario, the causal effect is indirectly mediated by third variables; for example, when *y*_*t−*2_ affects *z*_*t−*1_, which affects *x*_*t*_, we find the causal effect of *y*_*t−*2_ on *x*_*t*_. In the second scenario, target variables are strongly driven by external forces, which are never embedded by time-delayed variables. In this case, an external force *z*_*t−*2_ affects both *y*_*t−*1_ and *x*_*t*_, resulting in the spurious detection of a causal effect of *y*_*t−*1_ on *x*_*t*_. The misidentifications of causality are ameliorated by conducting the causality test conditional on third variables. In the following section, we introduce the conditional UIC test based on previous studies of TE [9].

### A. Conditional UIC

The measure of causal influence conditional on third variables, ***z*** = {*z*_*t*_}, is defined as follows:

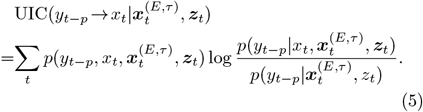

The difference from unconditional UIC defined by Eq. 4 is that the conditional probabilities include *z*_*t*_ as conditional variables. By conducting two causal tests:

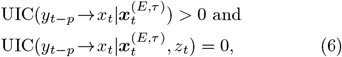

we can examine whether the causal influence of *y*_*t−p*_ on *x*_*t*_ is indirectly mediated by *z*_*t*_ or not.

### B. Applications

First, we applied the conditional UIC test to a four-species food chain model to demonstrate the identification of direct and indirect effects [12]. In the food chain model, one species directly affects another species at time-lag 1 along a transitive causal chain (Fig. 2; Supplemental Material S5). Using conditional UIC tests expressed by conditions (6), we successfully detected the direct causal effects at time lag 1 (red circle in Fig. 2; *P <* 0.05 using a surrogate-based significance test). However, when using only unconditional UIC tests, we detected many indirect effects (blue circle in Fig. 2) in addition to direct effects.

**FIG. 2.**
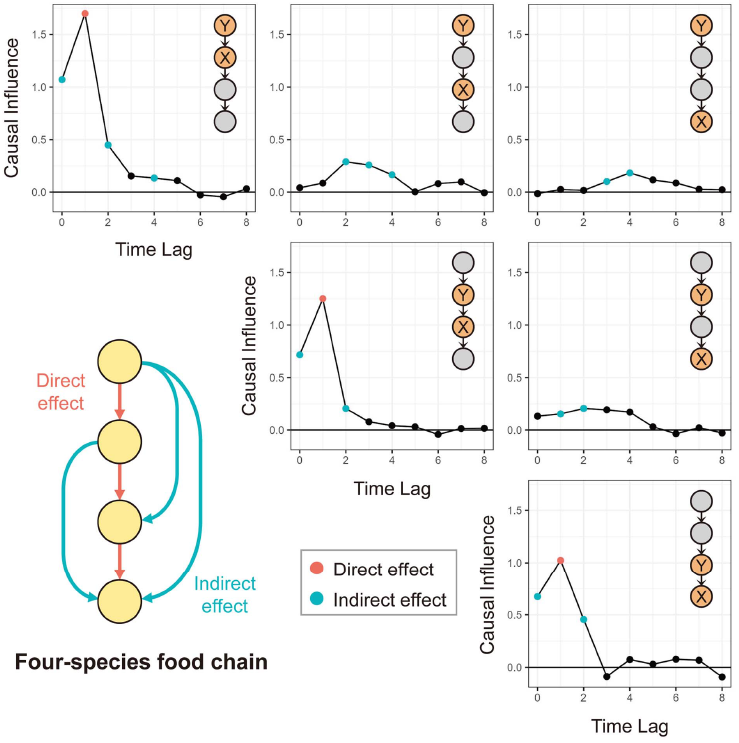
Identification of direct and indirect effects based on conditional UIC in a four-species food chain model. In this model, species directly affect their prey at time lag 1. The causality tests were performed for species pairs with direct effects (diagonal panels) and with indirect effects (off-diagonal panels). For each species pair, species X and Y were used as effect and cause variables, respectively. Red and blue circles indicate direct and indirect effects detected by significance tests, while black circles indicate no causal effects. Causal influences (vertical axes) are estimated by unconditional UIC (also see Supplemental Material Fig. S1).

Next, we applied the conditional UIC test to bacterial DNA time-series data collected from an experimental rice field [18]. These time-series data were obtained by quantifying DNA copy numbers for over 1000 taxa from irrigated water in experimental rice plots. In this application, we chose 20 abundant bacterial taxa with sufficient temporal fluctuations (see Supplemental Material S6 for more details). Conditional UIC detected 16 direct interactions and 28 indirect interactions among these bacterial taxa. To investigate the effects of identifying direct and indirect interactions on dynamical properties, we further estimated the signs and strengths of these interactions by S-map [19, 20]. Then, we evaluated the dynamic stability of the system [21] in two situations: (1) the system is incorrectly assumed to be driven by both direct and indirect interactions (Fig. 3a) and (2) the system is correctly assumed to be driven by only direct interactions (Fig. 3b). In this analysis, the sign was reversed for 4 of 16 direct interactions due to a failure to distinguish between indirect and direct interactions. These interactions had weak (i.e., near-zero) strengths, and their estimates may be influenced by other strong interactions. Using the conditional UIC, the system was identified as more stable because the reconstructed network eliminated suspicious indirect interactions (lower panels in Fig. 3), suggesting that the time-series analyses that do not distinguish between direct and indirect effects will misestimate dynamical properties such as system stability.

**FIG. 3.**
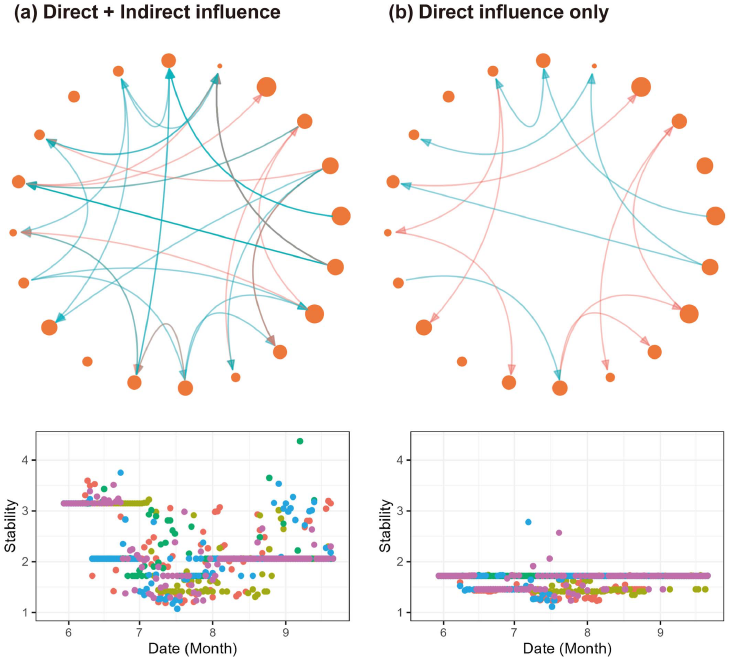
Application of conditional UIC to empirical bacterial community data. (a) Unconditional UIC detects 44 interactions between 20 Proteobacteria taxa. Each node size indicates the abundance of each bacterial DNA. Blue and red arrows represent positive and negative interactions estimated by S-map, respectively. The dynamic stability estimated from these interactions exhibits large temporal fluctuation. Colors in the scatter plots indicate different experimental plots. (b) Conditional UIC finds only 16 direct interactions among 44 interactions. The dynamic stability indicates that the system is more stable when estimated from only direct interactions.

## IV. DISCUSSION

In this paper, we develop a unified framework for non-parametric causal tests based on information-theoretic formulations and propose a novel causality test (i.e., UIC), establishing the theoretical connection between TE and CCM. Numerical experiments reveal that UIC shows higher robustness to noise in time-series data and have lower data requirements than TE and CCM (Fig. 1). Importantly, UIC outperforms TE and CCM in both linear and nonlinear dynamical systems. These results support the real-world applications of UIC because we do not know *a priori* whether TE or CCM is an appropriate causality test in weakly nonlinear systems.

In addition to its high performance, our method has two key advantages over conventional approaches. First, UIC has access to previous theoretical advances in both information theory [6, 10] and dynamical systems theory [22, 23]. As such, our framework facilitates the development of causality tests in an integrative way, with the help of both theories. Second, our method enables efficient computations of statistical significance. Specifically, we can reduce the computational requirements for searching nearest neighbors for state space reconstruction, which is the greatest computational challenge in nonparametric causality tests (Supplemental Material S3). Because recent studies have focused on large systems with thousands of components or more [18], efficient computation is important for real-world applications.

Interestingly, despite the mathematical equivalence of their definitions, TE and UIC show distinct statistical behavior. This arises from the following three characteristics of the nonparametric causality tests we studied: (1) the use of time-delayed variables for state space re-construction, (2) the need to search nearest neighbors to obtain nonparametric estimators, and (3) the need to separately calculate two (conditional) probabilities and their ratio to obtain causal influence. In contrast to (1) and (2), the last characteristic may depend on the algorithm used in nonparametric causality tests. For example, the Frenzel-Pompe algorithm [9] gives an identical value for causal influence for both TE and UIC. Although comparing computational algorithms is beyond the scope of our study, we stress the caveat that TE and UIC might have similar statistical behavior when implemented using other computational algorithms.

To understand complex systems in the real world, it is critical to identify interactions among components. For example, under anthropogenic stress, ecosystems sometimes exhibit unexpected regime shifts, including decreases in net primary production [24, 25] and mass bleaching of corals [26], due to transitions in dynamical structure emergent from interacting components [27]. With improved ecosystem monitoring techniques [28], nonparametric causality detection allows us to capture such critical transitions, providing a basis for preventing unexpected regime shifts and supporting ecosystem management to maintain biodiversity and ecosystem functions. Our method can improve the application of causality detection to real-world complex systems, thereby contributing to human well-being in a changing world.

## ACKNOWLEDGMENTS

This study was supported financially by JSPS KAKENHI grants (nos. 18K14801, 21K15171, 19H05641 and 21H05312).

