## Supplementary Materials for "Unified understanding of nonparametric causality detection in time series"

**Table of Contents**

S1. Transfer Entropy and Convergent Cross Mapping

S2. Unified Information-theoretic Causality

S3. Significance Test Based on the Surrogate Method

S4. Algorithm

S5. Setting of Numerical Experiments

S6. Empirical Microbial Community Data

Reference, Table S1, Figure S1, Figure S2

### S1. Transfer Entropy (TE) and Convergent Cross Mapping (CCM)

We provide the definitions of transfer entropy (Schreiber 2000) and convergence cross mapping (Sugihara *et al.* 2012) from the original papers and propose an information-theoretic definition of CCM. In this section, we consider the system with $\boldsymbol{x}$ as the effect variable and $\boldsymbol{y}$ as the cause variable. $\boldsymbol{x}_t^{(E,\tau)} = \{x_{t-\tau}, x_{t-2\tau} \dots, x_{t-E\tau}\}$ was used as a time-lagged vector in which the effect variable is embedded with the number of dimensions ($E > 0$) and time-lag ($\tau > 0$).

| **Definition 1: Transfer Entropy** (Schreiber 2000) |
|---|
| Let $x_t$ be embedded by $y_{t-p}$ and $\boldsymbol{x}_t^{(E,\tau)}$ based on time-delay embedding theorem. $p > 0$ is the causal time delay. Transfer entropy is defined as follows: $$\mathrm{TE}\left(y_{t-p} \to x_t \middle| \boldsymbol{x}_t^{(E,\tau)}\right) = \sum_t p\left(y_{t-p}, x_t, \boldsymbol{x}_t^{(E,\tau)}\right) \log \frac{p\left(x_t \middle| y_{t-p}, \boldsymbol{x}_t^{(E,\tau)}\right)}{p\left(x_t \middle| \boldsymbol{x}_t^{(E,\tau)}\right)}.$$ The causal influence from $y_{t-p}$ to $x_t$ is tested by the following inequality: $$\mathrm{TE}\left(y_{t-p} \to x_t \middle| \boldsymbol{x}_t^{(E,\tau)}\right) > 0.$$ |

| **Definition 2: Convergent Cross Mapping** (Sugihara *et al.* 2012) |
|---|
| Let $x_t$ be embedded by $y_{t-p}$ and $\boldsymbol{x}_t^{(E,\tau)}$ based on time-delay embedding theorem. $p > 0$ is the causal time delay. Using cross mapping, the forecast performance is defined as follows: $$\mathcal{X}(\boldsymbol{L}, \boldsymbol{P}) = \sum_{t \in \boldsymbol{P}} p\left(y_{t-p}, x_t, \boldsymbol{x}_t^{(E,\tau)}\right) \log p(\boldsymbol{x_L}, \boldsymbol{y_L}) \frac{\int d\boldsymbol{W}\, p(y_{t-p} \mid \boldsymbol{y_L}, \boldsymbol{W})\, p\left(x_t, \boldsymbol{x}_t^{(E,\tau)} \middle| \boldsymbol{W}, \boldsymbol{x_L}\right) p(\boldsymbol{W} \mid \boldsymbol{x_L})}{\int d\boldsymbol{W}\, p\left(x_t, \boldsymbol{x}_t^{(E,\tau)} \middle| \boldsymbol{W}, \boldsymbol{x_L}\right) p(\boldsymbol{W} \mid \boldsymbol{x_L})},$$ $$= \sum_{t \in \boldsymbol{P}} p\left(y_{t-p}, x_t, \boldsymbol{x}_t^{(E,\tau)}\right) \log p(\boldsymbol{x_L}, \boldsymbol{y_L}) \int d\boldsymbol{W}\, p(y_{t-p} \mid \boldsymbol{y_L}, \boldsymbol{W})\, p\left(\boldsymbol{W} \middle| x_t, \boldsymbol{x}_t^{(E,\tau)}, \boldsymbol{x_L}\right),$$ where $\boldsymbol{L}$ and $\boldsymbol{P}$ are the sets of time indices for state space reconstruction (i.e., training) and prediction (i.e., cross-validation), respectively. $\boldsymbol{x_L} = \{x_t \mid t \in \boldsymbol{L}\}$ and $\boldsymbol{y_L} = \{y_t \mid t \in \boldsymbol{L}\}$. $\boldsymbol{W}$ denotes the model parameters. The causal influence from $y_{t-p}$ to $x_t$ is tested by the following inequality: $$\mathrm{CCM}\left(y_{t-p} \to x_t \middle| \boldsymbol{x}_{t-\tau}^{(E)}\right) = \mathcal{X}(\boldsymbol{L}, \boldsymbol{P}) - \mathcal{X}(\boldsymbol{L}_{\mathbf{min}}, \boldsymbol{P}) > 0,$$ where $\boldsymbol{L}_{\mathbf{min}}$ are the sets of library time indices at the minimum size. |

Given that the dependence between $\boldsymbol{x}$ and $\boldsymbol{y}$ is conditional on $\boldsymbol{W}$, that is,

$$p(y_{t-p}|\boldsymbol{y_L},\boldsymbol{W}) = p\left(y_{t-p}\middle|\boldsymbol{y_L},\boldsymbol{W},x_t,\boldsymbol{x}_t^{(E,\tau)},\boldsymbol{x_L}\right)$$

for cross mapping, the expectation of $\mathcal{X}(\boldsymbol{L},\boldsymbol{P})$ can be calculated as follows:

$$\begin{aligned}
&\mathbb{E}[\mathcal{X}(\boldsymbol{L},\boldsymbol{P})] \\
&= \mathbb{E}\left[\sum_{t\in\boldsymbol{P}} p\left(y_{t-p},x_t,\boldsymbol{x}_t^{(E,\tau)}\right)\log p(\boldsymbol{x_L},\boldsymbol{y_L})\frac{\int d\boldsymbol{W}\, p(y_{t-p}|\boldsymbol{y_L},\boldsymbol{W})\, p\left(x_t,\boldsymbol{x}_t^{(E,\tau)}\middle|\boldsymbol{W},\boldsymbol{x_L}\right)p(\boldsymbol{W}|\boldsymbol{x_L})}{\int d\boldsymbol{W}\, p\left(x_t,\boldsymbol{x}_t^{(E,\tau)}\middle|\boldsymbol{W},\boldsymbol{x_L}\right)p(\boldsymbol{W}|\boldsymbol{x_L})}\right] \\
&= \mathbb{E}\left[\sum_{t\in\boldsymbol{P}} p\left(y_{t-p},x_t,\boldsymbol{x}_t^{(E,\tau)}\right)\log p(\boldsymbol{x_L},\boldsymbol{y_L})\frac{\int d\boldsymbol{W}\, p(y_{t-p}|\boldsymbol{y_L},\boldsymbol{W})\, p\left(x_t,\boldsymbol{x}_t^{(E,\tau)}\middle|\boldsymbol{W},\boldsymbol{x_L}\right)p(\boldsymbol{W}|\boldsymbol{x_L})}{p\left(x_t,\boldsymbol{x}_t^{(E,\tau)}\middle|\boldsymbol{x_L}\right)}\right] \\
&= \mathbb{E}\left[\sum_{t\in\boldsymbol{P}} p\left(y_{t-p},x_t,\boldsymbol{x}_t^{(E,\tau)}\right)\log p(\boldsymbol{x_L},\boldsymbol{y_L})\int d\boldsymbol{W}\, p(y_{t-p}|\boldsymbol{y_L},\boldsymbol{W})\, p\left(\boldsymbol{W}\middle|x_t,\boldsymbol{x}_t^{(E,\tau)},\boldsymbol{x_L}\right)\right] \\
&= \mathbb{E}\left[\sum_{t\in\boldsymbol{P}} p\left(y_{t-p},x_t,\boldsymbol{x}_t^{(E,\tau)}\right)\log p(\boldsymbol{x_L},\boldsymbol{y_L})p\left(y_{t-p}\middle|x_t,\boldsymbol{x}_t^{(E,\tau)},\boldsymbol{y_L},\boldsymbol{x_L}\right)\right] \\
&= \mathbb{E}\left[\sum_{t\in\boldsymbol{P}} p\left(y_{t-p},x_t,\boldsymbol{x}_t^{(E,\tau)}\right)\log p\left(y_{t-p}\middle|x_t,\boldsymbol{x}_t^{(E,\tau)}\right)\right].
\end{aligned}$$

Assuming $\boldsymbol{L}_{\mathbf{min}} \to \boldsymbol{\phi}$ (i.e., empty set), we can also calculate the expectation of $\mathcal{X}(\boldsymbol{L}_{\mathbf{min}},\boldsymbol{P})$.

$$\begin{aligned}
&\mathbb{E}[\mathcal{X}(\boldsymbol{\phi},\boldsymbol{P})] \\
&= \mathbb{E}\left[\sum_{t\in\boldsymbol{P}} p\left(y_{t-p},x_t,\boldsymbol{x}_t^{(E,\tau)}\right)\log\frac{\int d\boldsymbol{W}\, p(y_{t-p}|\boldsymbol{W})\, p\left(x_t,\boldsymbol{x}_t^{(E,\tau)}\middle|\boldsymbol{W}\right)p(\boldsymbol{W})}{\int d\boldsymbol{W}\, p\left(x_t,\boldsymbol{x}_t^{(E,\tau)}\middle|\boldsymbol{W}\right)p(\boldsymbol{W})}\right] \\
&= \mathbb{E}\left[\sum_{t\in\boldsymbol{P}} p\left(y_{t-p},x_t,\boldsymbol{x}_t^{(E,\tau)}\right)\log\frac{\int d\boldsymbol{W}\, p(y_{t-p}|\boldsymbol{W})p(\boldsymbol{W})}{\int d\boldsymbol{W}\, p(\boldsymbol{W})}\right] \\
&= \mathbb{E}\left[\sum_{t\in\boldsymbol{P}} p\left(y_{t-p},x_t,\boldsymbol{x}_t^{(E,\tau)}\right)\log p(y_{t-p})\right]
\end{aligned}$$

Thus, we obtain the difference in these expectations as follows:

$$\mathbb{E}[\mathcal{X}(\boldsymbol{L},\boldsymbol{P}) - \mathcal{X}(\boldsymbol{\phi},\boldsymbol{P})] = \mathbb{E}\left[\sum_{t} p\left(y_{t-p},x_t,\boldsymbol{x}_t^{(E,\tau)}\right)\log\frac{p\left(y_{t-p}\middle|x_t,\boldsymbol{x}_t^{(E,\tau)}\right)}{p(y_{t-p})}\right],$$

indicating that as the library (i.e., training) data size approaches zero, we can derive the information-theoretic definition of CCM.

**Definition 3: CCM based on Information Theory**

Let $x_t$ be embedded by $y_{t-p}$ and $\boldsymbol{x}_t^{(E,\tau)}$ based on time-delay embedding theorem. $p > 0$ is the causal time delay. Based on information theory, CCM is redefined as follows:

$$\mathrm{CCM}^{\mathrm{IT}}\left(y_{t-p} \to x_t \middle| \boldsymbol{x}_t^{(E,\tau)}\right) = \sum_t p\left(y_{t-p}, x_t, \boldsymbol{x}_t^{(E,\tau)}\right) \log \frac{p\left(y_{t-p} \middle| x_t, \boldsymbol{x}_t^{(E,\tau)}\right)}{p\left(y_{t-p}\right)}.$$

### S2. Unified Information-theoretic Causality

In this section, we define a novel causality measure, Unified Information-theoretic Causality (UIC) and its generalization with conditional variables (Frenzel & Pompe 2007).

**Definition 4: Unified Information-theoretic Causality**

Let $x_t$ be embedded by $y_{t-p}$ and $\boldsymbol{x}_t^{(E,\tau)}$ based on time-delay embedding theorem. $p > 0$ is the causal time delay. Unified information-theoretic causality is defined as follows:

$$\mathrm{UIC}\left(y_{t-p} \to x_t \middle| \boldsymbol{x}_{t-\tau}^{(E)}\right) = \sum_t p\left(y_{t-p}, x_t, \boldsymbol{x}_t^{(E,\tau)}\right) \log \frac{p\left(y_{t-p} \middle| x_t, \boldsymbol{x}_t^{(E,\tau)}\right)}{p\left(y_{t-p} \middle| \boldsymbol{x}_t^{(E,\tau)}\right)}.$$

The causal influence from $y_{t-p}$ to $x_t$ is tested by the following inequality:

$$\mathrm{UIC}\left(y_{t-p} \to x_t \middle| \boldsymbol{x}_t^{(E,\tau)}\right) > 0.$$

**Definition 5: Conditional Unified Information-theoretic Causality**

Let $x_t$ be embedded by $y_{t-p}$, $\boldsymbol{x}_t^{(E,\tau)}$ and $\boldsymbol{z}_t$ based on time-delay embedding theorem. $p > 0$ is the causal time delay. Conditional unified information-theoretic causality is defined as follows:

$$\mathrm{UIC}\left(y_{t-p} \to x_t \middle| \boldsymbol{x}_{t-\tau}^{(E)}, \boldsymbol{z}_t\right) = \sum_t p\left(y_{t-p}, x_t, \boldsymbol{x}_t^{(E,\tau)}, \boldsymbol{z}_t\right) \log \frac{p\left(y_{t-p} \middle| x_t, \boldsymbol{x}_t^{(E,\tau)}, \boldsymbol{z}_t\right)}{p\left(y_{t-p} \middle| \boldsymbol{x}_t^{(E,\tau)}, \boldsymbol{z}_t\right)}$$

The causal influence from $y_{t-p}$ to $x_t$ not mediated by $\boldsymbol{z}_t$ is tested by the following inequality:

$$\mathrm{UIC}\left(y_{t-p} \to x_t \middle| \boldsymbol{x}_t^{(E,\tau)}, \boldsymbol{z}_t\right) > 0.$$

From Definitions 4 and 5, the indirect causal influence from $y_{t-p}$ to $x_t$ mediated by $\boldsymbol{z}_t$ needs to satisfy the following two conditions:

$$\mathrm{UIC}\left(y_{t-p} \to x_t \middle| \boldsymbol{x}_{t-\tau}^{(E)}\right) > 0 \text{ and } \mathrm{UIC}\left(y_{t-p} \to x_t \middle| \boldsymbol{x}_{t-\tau}^{(E)}, \boldsymbol{z}_t\right) = 0.$$

**S3. Significance Test Based on the Surrogate Method**

Based on the surrogate method, we derive an efficient significance test for UIC. The surrogate methods requires the generation of data that artificially exclude the target causal influence, with all else being equal (surrogate data). As such, the conditions to generate surrogate data should satisfy the following dependence (Runge *et al.* 2018):

$$p\left(y_{t-p} \middle| x_t, \boldsymbol{x}_t^{(E,\tau)}\right) = p\left(y_{t-p} \middle| \boldsymbol{x}_t^{(E,\tau)}\right).$$

Thus, we obtain the following equation:

$$\sum_t p\left(y_{t-p}, x_t, \boldsymbol{x}_t^{(E,\tau)}\right) \log \frac{p\left(y_{t-p} \middle| x_t, \boldsymbol{x}_t^{(E,\tau)}\right)}{p\left(y_{t-p} \middle| \boldsymbol{x}_t^{(E,\tau)}\right)}$$

$$= \sum_t p\left(y_{t-p} \middle| \boldsymbol{x}_t^{(E,\tau)}\right) p\left(x_t, \boldsymbol{x}_t^{(E,\tau)}\right) \log \frac{p\left(y_{t-p} \middle| x_t, \boldsymbol{x}_t^{(E,\tau)}\right)}{p\left(y_{t-p} \middle| \boldsymbol{x}_t^{(E,\tau)}\right)},$$

indicating that the surrogate data for $y_{t-p}$ should be generated conditional on $\{x_{t-\tau}, \dots, x_{t-E\tau}\}$ for UIC. It is noteworthy that $y_{t-p}$ is not included in the set of conditional variables for state space reconstruction. This allows us to implement efficient significance tests because the highest computational cost of nonparametric causality tests is to search nearest neighbors for state space reconstruction. In this study, we simply implement surrogate methods based on cross mapping (cross-map surrogate method).

**S4. Algorithm**

The UIC algorithm to detect causality from observed time-series data is based on cross mapping (Sugihara *et al.* 2012), with some modifications. Cross mapping from $\boldsymbol{x}$ to $\boldsymbol{y}$ is interpreted as a k-nearest neighbor (k-NN) regression with time-delayed variables ($\boldsymbol{x}$ is explanatory variables and $\boldsymbol{y}$ is response variables). Let $\boldsymbol{\omega}$ be time-delayed variables of $\boldsymbol{x}$. The cross mapping uses:

$$w_{t,i} = \exp\left(-\frac{\|\boldsymbol{\omega}_t - \boldsymbol{\omega}_i\|}{\left\|\boldsymbol{\omega}_t - \boldsymbol{\omega}_{nn(t)}\right\|}\right)$$

as the weights of k-NN regression. $\|\cdot\|$ is the distance between time-delayed variables with different time indices. $nn(t)$ is the time index of the nearest neighbor of $\boldsymbol{\omega}_t$. Then, the predicted value of $y_t$ is:

$$\hat{y}_t \mid \boldsymbol{\omega} = \sum_{i \in NN_k(t)} w_{t,i} y_i,$$

where $NN_k(t) \in L$ is the set of $k$ nearest neighbors of $\boldsymbol{\omega}_t$. In the original paper (Sugihara *et al.* 2012), the mean squared error between observed and predicted values was used to evaluate model performance:

$$\mathrm{MSE}_{\text{naïve}}(\boldsymbol{y}|\boldsymbol{\omega}) = \frac{1}{n_p}\sum_{t \in \boldsymbol{P}}\left\{y_{t-p} - (\hat{y}_{t-p} \mid \boldsymbol{\omega})\right\}^2.$$

However, $\mathrm{MSE}_{\text{naïve}}$ is a biased variance estimator of $y_t|\boldsymbol{w}$ for a small number of the nearest neighbors. Our algorithm used an alternative unbiased variance estimator:

$$\mathrm{MSE}(\boldsymbol{y}|\boldsymbol{\omega}) = \frac{1}{n_p}\sum_{t \in \boldsymbol{P}}\left(1 + \frac{1}{k_{\text{eff}}(t)}\right)^{-1}\left\{y_{t-p} - (\hat{y}_{t-p} \mid \boldsymbol{\omega})\right\}^2,$$

where $k_{\text{eff}}(t) = 1/\sum_{i \in NN_k(t)} w_{t,i}^2$ is the effective number of nearest neighbors. The derivation of $\mathrm{MSE}(\boldsymbol{y}|\boldsymbol{\omega})$ is provided at the end of this section.

To express the causal influence of UIC with mean squared errors, we assume that the errors, $\varepsilon_t = y_t - (\hat{y}_t \mid \boldsymbol{\omega})$, are independent identically distributed (iid) normal random variables with a constant variance for all time indices. If this assumption is satisfied, then we derive the estimated causal influence from $y_{t-p}$ to $x_t$:

$$\widehat{\mathrm{UIC}}\left(y_{t-p} \to x_t \middle| \boldsymbol{x}_{t-\tau}^{(E)}\right) = -\frac{1}{2}\left\{\log\left(\mathrm{MSE}\left(\boldsymbol{y}\middle|x_t, \boldsymbol{x}_{t-\tau}^{(E)}\right)\right) - \log\left(\mathrm{MSE}\left(\boldsymbol{y}\middle|\boldsymbol{x}_{t-\tau}^{(E)}\right)\right)\right\}.$$

The expectation of this estimator coincides with that of theoretical value,

$$\mathbb{E}\left[\widehat{\mathrm{UIC}}\left(y_{t-p} \to x_t \middle| \boldsymbol{x}_{t-\tau}^{(E)}\right)\right] = \mathbb{E}\left[\mathrm{UIC}\left(y_{t-p} \to x_t \middle| \boldsymbol{x}_{t-\tau}^{(E)}\right)\right]$$

because under the assumption of iid normal random variables,

$$\mathbb{E}\left[\sum_{t} p(y_{t-p}, \boldsymbol{\omega}) \log p(y_{t-p}|x_t, \boldsymbol{\omega})\right] = -\frac{1}{2}\left\{\mathbb{E}[\log(\mathrm{MSE}(\boldsymbol{y}|\boldsymbol{\omega}))] + \log(2\pi) + \frac{n_l}{n_p}\right\}.$$

The UIC algorithm was implemented as an R package (https://github.com/yutakaos/rUIC). Finally, we provide the derivation of $\mathrm{MSE}(\boldsymbol{y}|\boldsymbol{\omega})$. When $1 \ll n_l$, the following equations are obtained:

$$\mathbb{E}[y_t] = \mathbb{E}\left[\sum_{i \in NN_k(t)} w_{t,i} y_t\right] \text{ and } \mathbb{E}[y_t] \approx \mathbb{E}\left[\sum_{i \in NN_k(t)} w_{t,i}^2 y_t \Big/ \sum_{i \in NN_k(t)} w_{t,i}^2\right].$$

Thus,

$$\mathbb{E}\left[\left\{y_t - \sum_{i} w_{t,i} y_i\right\}^2\right] = \left(1 + \sum_{i} w_{t,i}^2\right)(\mathbb{E}[y_t^2] - \mathbb{E}[y_t]^2).$$

By definition, $\mathrm{MSE}(y_t) = \mathbb{E}[(y_t - \mathbb{E}[y_t])^2] = \mathbb{E}[y_t^2] - \mathbb{E}[y_t]^2$, we can obtain an unbiased variance estimator:

$$\mathrm{MSE}(\boldsymbol{y}|\boldsymbol{\omega}) = \frac{1}{n_p}\sum_{t \in \boldsymbol{P}}\left(1 + \sum_{i} w_{t,i}^2\right)^{-1}\left\{y_t - \sum_{i} w_{t,i} y_i\right\}^2.$$

**S5. Numerical Experiments**

To compare the performance of causality tests, we generated synthetic time-series data from two dynamical systems with noise: (1) a nonlinear two-component logistic model:

$$x_t = x_{t-1}(3.5 - 3.5\, x_{t-1} - \beta\, y_{t-1}), \qquad X_t = x_t + \varepsilon_t^X$$

$$y_t = y_{t-1}(3.8 - 3.8\, y_{t-1} - 0.02\, x_{t-1}), \qquad Y_t = y_t + \varepsilon_t^Y$$

and (2) a linear two-component VAR model:

$$x_t = 0.1\, x_{t-1} + \beta\, y_{t-1} + \varepsilon_t^X, \qquad X_t = x_t$$

$$y_t = 0.1\, y_{t-1} + \varepsilon_t^Y, \qquad Y_t = y_t$$

where the observed time-series $\boldsymbol{X} = \{X_t\}$ and $\boldsymbol{Y} = \{Y_t\}$ were obtained from true model dynamics of $\boldsymbol{x} = \{x_t\}$ and $\boldsymbol{y} = \{y_t\}$, respectively. The causal influences are controlled by the parameter $\beta$. Then, we conducted two numerical experiments. In the first experiment, we performed model simulations with different noise levels for effect variables ($\varepsilon_t^X$) and cause variables ($\varepsilon_t^Y$). We used $\mathrm{sd}(\boldsymbol{\varepsilon}^{\boldsymbol{X}})/\mathrm{sd}(\boldsymbol{x}) =$ 0.01 or 0.5 and $\mathrm{sd}(\boldsymbol{\varepsilon}^{\boldsymbol{Y}})/\mathrm{sd}(\boldsymbol{y}) =$ 0.01 or 0.5 for the logistic model and $\mathrm{sd}(\boldsymbol{\varepsilon}^{\boldsymbol{X}}) =$ 1.0 or 2.0 and $\mathrm{sd}(\boldsymbol{\varepsilon}^{\boldsymbol{Y}}) =$ 1.0 or 2.0 for the VAR model (upper panels in Fig. 1 in main text). In the second experiment, we varied the number of time points in time-series data, $T$ = 10, 20, …, 300 at the same noise level (lower panels in Fig. 1 in main text). Each experiment was repeated 1000 times without causality ($\beta$ = 0) and 1000 times with causality ($\beta$ > 0) and the performance of causality tests was evaluated in terms of the area under the receiver-operator characteristic curve (AUC). In the experiments with causality, we used $\beta$ = 0.1 for the logistic model and $\beta$ = 0.5 for the VAR model.

To demonstrate the identification of direct and indirect effects, we also conducted numerical experiments using a four-component-chain logistic model (Fig. S1):

$$x_t = x_{t-1}(3.9 - 3.9\, x_{t-1}),$$

$$y_t = y_{t-1}(3.6 - 3.6\, y_{t-1} - 0.40\, x_{t-1}),$$

$$z_t = z_{t-1}(3.6 - 3.6\, z_{t-1} - 0.40\, y_{t-1}),$$

$$w_t = w_{t-1}(3.8 - 3.8\, w_{t-1} - 0.35\, z_{t-1}),$$

where $\boldsymbol{x} = \{x_t\}$, $\boldsymbol{y} = \{y_t\}$, $\boldsymbol{z} = \{z_t\}$ and $\boldsymbol{w} = \{w_t\}$ were used as observed time-series without noise.

All analyses were conducted using R version 4.2.2 (R core team 2022). We utilized the *RTransferEntropy* package (version 0.2.14, Behrendt *et al.* 2019) and *rEDM* package (version 0.7.4, Ye *et al.* 2023) for TE and CCM, respectively. We implemented UIC algorithms (see Appendix S4) in the *rUIC* package (version 0.9.0), obtained from https://github.com/yutakaos/rUIC. The *pROC* package (version 1.18.0, Robin *et al.* 2011) also was used to compute AUC. Codes has been deposited in GitHub (https://github.com/yutakaos/archives/tree /master/analysis/2023_uic).

### S6. Empirical microbial community data

The rice growth experiments were conducted in five experimental plots, started on 23 May 2017 and finished on 22 September 2017 (Ushio 2022; https://doi.org/10.5281/zenodo.6510542). Irrigated water samples were collected from each plot every day during the experimental period, and the environmental DNA copy numbers for 1197 taxa in water samples were quantified. Among 1197 taxa, we chose 20 abundant bacteria with sufficient temporal fluctuations (belonging to the phylum Proteobacteria; Table S1) to examine causal influences. Proteobacteria was the most abundant and most diverse phylum in this experiment, and these 20 Proteobacteria were detected frequently in July. The DNA dynamics of these proteobacteria are shown in Fig. S2. The DNA copies were standardized by the maximum number of DNA copies of each taxon (i.e., scaled from 0 to 1). Using all time-series data for 20 Proteobacteria, we conducted causality tests based on unconditional and conditional UIC and estimated the sign and strength of the detected causal relations by S-map (Sugihara 1994). Then, we evaluated the dynamic stability of the bacterial community assuming the system is driven by the detected causal relations (Ushio *et al.* 2018).

**Table S1 |** Proteobacteria taxa used in causality tests. Taxa were assigned by the lowest common ancestor algorithm. For details, see Ushio (2022).

| Taxa ID | Phylum – Class – Order – Family – Genus |
|---|---|
| Prok_Taxa00004 | Proteobacteria – Betaproteobacteria – Burkholderiales |
| Prok_Taxa00006 | Proteobacteria – Betaproteobacteria – Burkholderiales – Burkholderiaceae – *Polynucleobacter* |
| Prok_Taxa00007 | Proteobacteria – Alphaproteobacteria – Rhizobiales |
| Prok_Taxa00009 | Proteobacteria – Alphaproteobacteria – Sphingomonadales – Sphingomonadaceae |
| Prok_Taxa00016 | Proteobacteria – Alphaproteobacteria – Rhizobiales – Hyphomicrobiaceae |
| Prok_Taxa00028 | Proteobacteria – Alphaproteobacteria – Rhodospirillales – Rhodospirillaceae – *Azospirillum* |
| Prok_Taxa00029 | Proteobacteria – Alphaproteobacteria – Sphingomonadales – Sphingomonadaceae – *Novosphingobium* |
| Prok_Taxa00034 | Proteobacteria |
| Prok_Taxa00038 | Proteobacteria – Betaproteobacteria – Burkholderiales – Comamonadaceae |
| Prok_Taxa00041 | Proteobacteria – Betaproteobacteria – Burkholderiales |
| Prok_Taxa00121 | Proteobacteria – Alphaproteobacteria – Sphingomonadales – Sphingomonadaceae |
| Prok_Taxa00146 | Proteobacteria – Alphaproteobacteria – Rhodospirillales – Rhodospirillaceae – *Lacibacterium* |
| Prok_Taxa00148 | Proteobacteria – Betaproteobacteria – Nitrosomonadales |
| Prok_Taxa00160 | Proteobacteria – Alphaproteobacteria – Rhodospirillales – Rhodospirillaceae – *Magnetospirillum* |
| Prok_Taxa00171 | Proteobacteria – Alphaproteobacteria |
| Prok_Taxa00201 | Proteobacteria – Gammaproteobacteria – Cellvibrionales – Cellvibrionaceae – *Cellvibrio* |
| Prok_Taxa00244 | Proteobacteria – Betaproteobacteria – Neisseriales – Chromobacteriaceae –Deefgea |
| Prok_Taxa00350 | Proteobacteria – Gammaproteobacteria |
| Euk_Taxa00117 | Proteobacteria – Alphaproteobacteria – Sphingomonadales –Sphingomonadaceae |
| Euk_Taxa00174 | Proteobacteria – Alphaproteobacteria – Rhizobiales – Hyphomicrobiaceae |

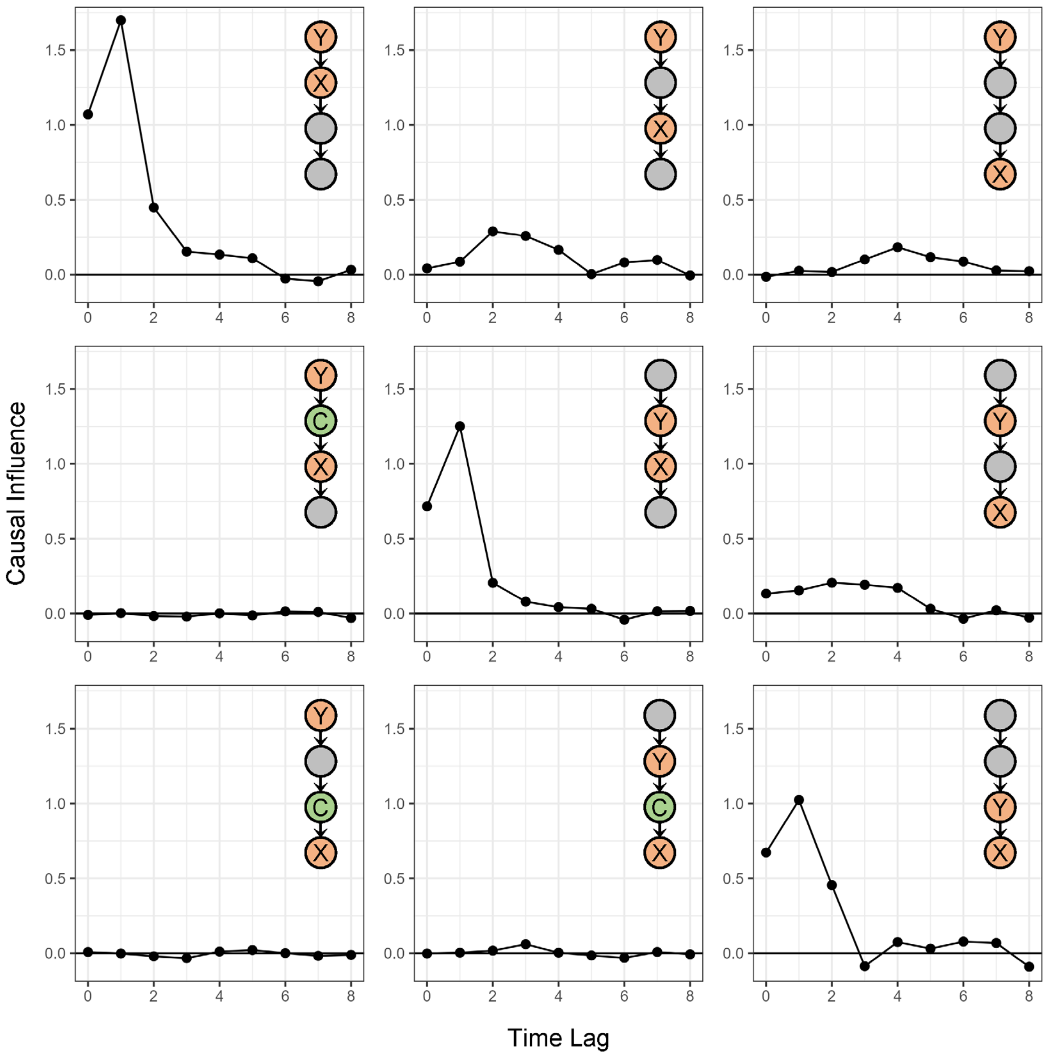


**Figure S1 |** Results of unconditional and conditional UIC tests in a four-species food chain model. For each test, X, Y and C were used as effect, cause and conditional variables, respectively.

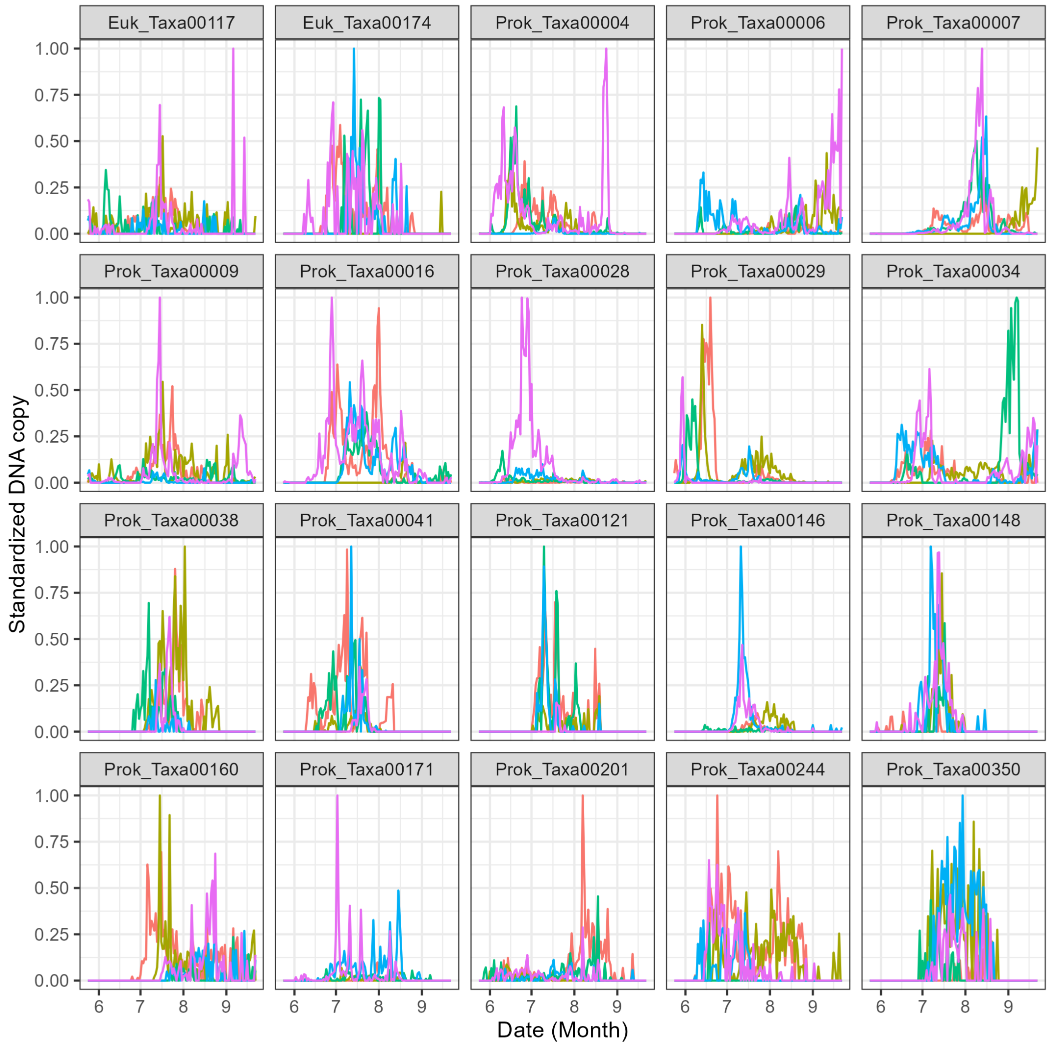


**Figure S2 |** Time-series data for DNA copy numbers of 20 Proteobacteria taxa. Time series were standardized by the maximum number of DNA copies for each taxon. Water samples were taken from five experimental plots. Different colors indicate different experimental plots. For details, see Ushio (2022).
